# Combined Inhibition of AKT and KIT Restores Expression of Programmed Cell Death 4 (PDCD4) in Gastrointestinal Stromal Tumor

**DOI:** 10.1101/2021.04.13.439680

**Authors:** Marya Kozinova, Shalina Joshi, Shuai Ye, Martin G. Belinsky, Dinara Sharipova, Jeffrey M. Farma, Sanjay Reddy, Samuel Litwin, Karthik Devarajan, Alex Rosa Campos, Yi Yu, Brian Schwartz, Margaret von Mehren, Lori Rink

**Author notes:** Correspondence to: Lori Rink, Ph.D., Molecular Therapeutics Program, Fox Chase Cancer Center, 333 Cottman Avenue, Philadelphia, PA 19111-2497.

## Abstract

The majority of gastrointestinal stromal tumor (GIST) patients develop resistance to the first-line KIT inhibitor, imatinib mesylate (IM), through acquisition of secondary mutations in *KIT* or bypass signaling pathway activation. AKT is a relevant target for inhibition, in addition to KIT, since the PI3K/AKT pathway is crucial for IM-resistant GIST survival. We evaluated the activity of a novel pan-AKT inhibitor, MK-4440 (formerly ARQ 751), as monotherapy and in combination with IM in GIST cell lines and preclinical models with varying IM sensitivities. Dual inhibition of KIT and AKT demonstrated significant synergistic effects in IM-sensitive and -resistant GIST cell lines. Proteomic analyses revealed upregulation of the tumor suppressor, PDCD4, in combination treated cells. Enhanced PDCD4 expression correlated to cell cycle arrest and cell death. *In vivo* studies revealed superior efficacy of MK-4440/IM combination in an IM-sensitive preclinical model of GIST compared with either single agent. The combination demonstrated limited efficacy in two IM-resistant models, including a GIST patient-derived xenograft model possessing an exon 9 *KIT* mutation. These studies provide strong rationale for further use of AKT inhibition in combination with IM in primary GIST; however, alternative agents will need to be tested in combination with AKT inhibition in the resistant setting.

## Introduction

Gastrointestinal stromal tumor (GIST) is the most common mesenchymal tumor of the gastrointestinal tract, with an annual worldwide incidence of 11 to 19.6 per million (1). The majority (85-90%) of GIST cases are caused by oncogenic mutations in the receptor tyrosine kinases (RTKs), KIT or PDGFRA, which result in constitutive activation of these receptors. The remaining GIST cases that lack mutations in these genes, typically gastric GIST, often lack expression of succinate dehydrogenase subunit B (SDHB) due to genetic or epigenetic deficiencies in the SDH complex of the respiratory chain (2, 3).

Surgery remains the first-line treatment for resectable GIST; however, some tumors are not able to be removed due to size, anatomic site, or in the metastatic setting. In addition, almost 30% of patients experience a recurrence of GIST within 5 years following surgery (4). Imatinib mesylate (IM), an RTK inhibitor targeting the mutant forms of KIT and PDGFRA, is the first-line therapy in the neoadjuvant setting for treatment of unresectable and metastatic GIST, to downstage the tumor prior to resection and in the adjuvant setting for patients with a high risk of recurrence for IM sensitive tumors (5-7). Treatment with IM leads to disease stabilization in more than 80% of patients; however, median time to tumor progression is approximately two years (8). The RTK inhibitors sunitinib (9) and regorafinib (10) are approved as second and third-line treatments, respectively, for the treatment of IM-refractory GIST, but only confer additional disease stabilization for less than 6 months. In 2019, avapritinib was approved for patients whose tumors have the IM-refractory PDGFRA exon 18 mutations (11) and more recently, in 2020, ripretinib (12, 13), a switch control kinase inhibitor with broad spectrum activity against RTK inhibitor resistant mutations, was approved for the treatment of GIST refractory to standard therapies.

Response to RTK inhibitors has been correlated to tumor genotype (14, 15). The majority of GIST harbor a mutation in exon 11 of KIT, encoding the juxtamembrane region of the protein, and are typically sensitive to IM. Patients with GIST that have mutations in KIT exon 9, encoding the extracellular domain of the RTK, typically require a higher dose of IM. GIST harboring primary *KIT* exon 13 or exon 17 mutations (encoding the ATP-binding site and activation-loop, respectively), and other exons occur in less than 5% of GIST cases, and are mostly resistant to IM (1, 16). Following a period of response to RTK therapy, the majority of tumors ultimately develop resistance, mainly through the acquisition of secondary RTK mutations and the activation of bypass RTK downstream pathways, such as the PI3K/AKT pathway (17, 18). While there are now five RTK inhibitors approved for treatment of GIST, there remains an unmet clinical need for treatment options for patients that have progressed on all approved lines of therapy. IM remains the agent with the greatest efficacy for the majority of GIST patients, as well as, the greatest tolerability. Therefore, preventing the development of acquired resistance to IM would be of benefit.

AKT activation has previously been shown to be a hallmark of IM resistance in GIST (19-22), Simultaneous targeting of AKT in addition to inhibiting KIT with IM has potential to be a promising strategy to prevent the development of secondary resistance to IM. Our group has previously shown significantly enhanced combination effects of IM with MK-2206, an oral pan-AKT inhibitor, in an IM-sensitive xenograft model *in vivo*, as measured by extended disease stabilization and improved survival (19). In this study, we evaluated the efficacy of a new, selective, allosteric pan-AKT inhibitor, MK-4440 (formerly ARQ 751) in both IM-sensitive and resistant GIST *in vitro* and *in vivo* models, including a GIST patient-derived xenograft (PDX) possessing an exon 9 *KIT* mutation. We report superior activity of MK-4440 in combination with IM in a panel of IM-sensitive and IM-resistant GIST cell lines. This activity is mediated by increased PDCD4 expression leading to increased apoptosis and cell cycle arrest. Furthermore, dual inhibition of KIT and AKT provide impressive disease stabilization in IM-sensitive GIST xenografts and trends toward stabilization in IM-resistant models *in vivo*.

## Material and Methods

### Cell lines, Patient derived xenograft model, Compounds, and Antibodies

The GIST-T1 tumor cell line possessing a heterozygous mutation in *KIT* exon 11, was kindly provided by Takahiro Taguchi (Kochi University, Kochi, Japan) (23). The GIST-T1/829 subline derived from parental GIST-T1 cells and possessing a secondary A829P kinase domain mutation, and the GIST430 tumor cell line possessing a primary *KIT* exon 11 deletion with a secondary mutation (V654A substitution) were all generously provided by Jonathan A. Fletcher (Dana Farber Cancer Institute, Boston, MA, USA). GIST-T1, GIST-T1/829 and GIST430 cells were grown as previously described (19). Cells were routinely monitored by Sanger sequencing to confirm their *KIT* mutation status and cell line identity. Imatinib mesylate (IM) (Gleevec™) was obtained from Selleckchem (Houston, TX, USA), dissolved in sterile PBS and stored at −20°C. MK-4440 (ARQ 751) was obtained from ArQule Inc., as solution dissolved in 0.1N Phosphoric Acid and stored at −20°C. All antibodies used in this study were purchased from Cell Signaling Technologies (Beverly, MA, USA), except p21 (Santa Cruz, CA, USA), and used according to the manufacturer’s instructions.

### Cell Proliferation/Viability Assay

To test *in vitro* drug sensitivity, tumor cells were plated in 96-well plates at optimal seeding densities in complete media and incubated overnight. Wells were then treated with varying doses of MK-4440 and/or IM. Cell proliferation and viability were measured at 72 hours post treatment using the CellTiter Blue Viability Assay (Promega, Fitchburg, WI, USA) as described previously (24). Assays were performed as three independent biological replicates, with minimum of six technical replicates in each treatment arm. Combination indexes of CI_LD50_ (GIST-T1) and CI_LD30_ (GIST430, GIST-T1/829) were quantified using the previously described approach (24).

### BrdU incorporation assay

BrdU incorporation assay was used to measure cell proliferation. GIST-T1 cells were labeled with BrdU for 4 hours and GIST430 cells were labeled for 20h, and then analyzed with BrdU-APC Flow Kit (BD Biosciences) as described previously (24).

### Preparation of Whole Cell Extract from Cells and Immunoblot Assays

The whole cell extracts (WCE) were prepared and evaluated by immunoblot as described previously (20).

### Protein Extraction and LC-MS/MS Analysis

GIST cell lines were treated for 20 hours with IM/MK-4440 combination at the following concentrations, 40nM/120nM (GIST-T1), 1uM/3uM (GIST-T1/829) or corresponding amount of DMSO (control) in three biological replicates.

Protein extracts were lysed with 8M urea 50 mM ammonium bicarbonate, and protein concentration was determined using a bicinchoninic acid (BCA) protein assay (Thermo Fisher Scientific, Waltham, MA, USA). Proteins were then digested, acidified with formic acid and subsequently desalted using AssayMap C18 cartridges (Agilent, Santa Clara, CA, USA) mounted on an Agilent AssayMap BRAVO liquid handling system. More detailed procedures on the preparation of proteins prior to liquid chromatography-tandem mass spectrometry (LC-MS/MS) analysis are described in the **Supplementary Methods S1**. Dried peptide fractions were reconstituted with 2% acetonitrile (ACN), 0.1% formic acid (FA) and analyzed by LC-MS/MS using a Proxeon EASY nanoLC system (Thermo Fisher Scientific) coupled to an Orbitrap Fusion Lumos mass spectrometer (Thermo Fisher Scientific). Peptides were separated using an analytical C18 Aurora column (75µm x 250 mm, 1.6µm particles; IonOpticks, Fitzroy, Australia) at a flow rate of 300nL/min using a 75-min gradient (80% ACN, 0.1% FA). The mass spectrometer was operated in positive data-dependent acquisition mode. MS1 spectra were measured in the Orbitrap with a resolution of 60 000, at accumulation gain control (AGC) target of 4e5 with maximum injection time of 50ms, and within a mass range from 375 to 1 500 m/z. The instrument was set to run in top speed mode with 1-second cycles for the survey and the MS/MS scans. After a survey scan, tandem MS was performed on the most abundant precursors with charge state between +2 and +7 by isolating them in the quadrupole with an isolation window of 0.7 m/z. Precursors were fragmented with higher-energy collisional dissociation (HCD) with normalized collision energy of 30% and the resulting fragments were detected in the ion trap in rapid scan mode at AGC of 1e4 and maximum injection time of 35ms. The dynamic exclusion was set to 20sec with a 10 ppm mass tolerance around the precursor. All mass spectrometry proteomics data including the raw data (“.RAW”) and search result files from MaxQuant have been deposited to ProteomeXchange Consortium via MASSIVE partner repository with the dataset identifier PXD023717.

### Data Processing and Analysis

All mass spectra were analyzed with MaxQuant software version 1.5.5.1. MS/MS spectra were searched against the curated (Swiss-Prot) Homo sapiens Uniprot protein sequence database (downloaded in January 2019) and GPM cRAP sequences (commonly known protein contaminants). Log2-transformed LFQ intensities for all 5 407 detected proteins in four treatment subgroups (GIST-T1, GIST-T1+IM/MK-4440, GIST-T1/829, GIST-T1/829+IM/ MK-4440) in three biological replicates, including the potential contaminants, zero intensity proteins, as well as single-peptide identified proteins are contained in **Data file S1**. Common contaminants and proteins identified by site only were removed and only proteins identified by >1 unique peptides (4 872 proteins) were qualified for subsequent analysis using Perseus software (1.6.10.50) (25). Only proteins identified in all three biological replicates in at least one experimental condition (control or under IM/ MK-4440treatment) were further considered for statistical analysis (totally 3 120 proteins in GIST-T1/829 cell line subgroup and 2 837 proteins in GIST-T1 subgroup). In order to save normal distribution of the Log_2_ transformed intensity histogram and to simulate signals from low abundant proteins, the filtered LFQ data were subjected to imputation of missing values performed from the normal distribution (width 0.3 and down shift 1.8) (26). After normalization, Pearson’s Correlation was used to identify potential outliers, as a result there were no replicates removed from the analysis.

Two-sided Student’s t-tests were performed to identify significantly changing proteins among control and IM/ MK-4440 treated cells. *P-*values were adjusted for using permutation-based false-discovery rate (FDR = 0.05, s0 = 2, and 250 randomizations). **Data file S2** contains lists proteins qualified for the analysis, as well as significance test results. Volcano plots depicting Log2 fold change differences in protein abundance following IM/MK-4440 treatment was generated with Perseus for each cell line. Proteins significantly up- and downregulated upon IM/ MK-4440 treatment were compared between GIST-T1 and GIST-T1/829 cell lines with Venny 2.1 plotter (https://bioinfogp.cnb.csic.es/tools/venny/index.html).

### Establishment of GIST Patient Derived Xenografts

All studies involving animals followed procedures approved by the FCCC Institutional Animal Care and Use Committee. After obtaining an informed consent (FCCC IRB 03-848), GIST tumor tissues from a patient were freshly collected from the pathology department. Unfixed human tissue was transported using BSL-2 practices in chilled isotonic solution supplemented with antibiotics. NSG mice were anesthetized with inhaled isofluorane (1-3% in oxygen) and injection site was disinfected. Tissue was cut on ice into 2-3mm fragments, then mixed 1:1 (v/v) in full DMEM media/Matrigel (Corning Inc., Corning, NY, USA), minced to pass through a 16-gauge needle and implanted subcutaneously in both flanks of NSG mice (NSG-SGM3, obtained from the FCCC breeding colony) at a final volume of 200ml per injection. Additional fragments of tumor tissue were snap-frozen for DNA extraction or fixed in 10% formalin and processed to paraffin blocks for histological evaluation. For serial transplantation, tumors were extracted from euthanized tumor-bearing animals, minced under sterile conditions as described above, and injected in successive NSG mice. Once a tumor was established (PDX9.1), typical GIST histological features (KIT expression, cellular morphology) and genetic identity using DNA sequencing (alteration in exon 9 of KIT, p.A502_Y503dup) was confirmed over multiple passages and tumor from passage 8 was ultimately used for the drug administration study.

### GIST Xenografts and Drug Administration

GIST-T1 and GIST-430 cells were washed and subsequently resuspended in phosphate-buffered saline (PBS) at a density of 3 × 10^6^ cells/100 μl and 1 × 10^6^ cells/100 μl respectively, cells were mixed thoroughly with Corning Matrigel Matrix in 1:1 (v/v) ratio. The cell suspensions, as well as the previously described PDX9.1 tissue suspensions, were injected subcutaneously into the right flanks of SCID and NSG mice, respectively. Growth of established tumor xenografts was monitored at least twice a week by vernier caliper measurement of the smallest and largest tumor’s diameters and tumor volume was calculated using the formula: tumor volume (mm^3^) = smallest diameter^2^ × largest diameter × (*π* / 6). When tumors reached approximately 200 mm^3^, mice were randomized into four treatment arms: arm 1, vehicle (5 days/week, oral); arm 2, IM at 50 mg/kg (5 days/week, oral); arm 3, MK-4440 at 25mg/kg (4 days/week, oral); and arm 4, IM and MK-4440 at monotherapy doses. Treatment was continued until the tumors exceeded >10% of their body weight or the animals demonstrated distress or weight loss >10% as per the local IACUC guidelines. Tumors were harvested and frozen in dry ice.

### Tumor growth modeling

Tumor volume was measured for every mouse in each of the four treatment arms (vehicle, IM, MK-4440, and combination) in all three GIST models at a total of 20 distinct time points, from baseline (day 0) until study conclusion (day 67) for GIST-T1 xenografts, 20 distinct time points (until day 66) for PDX9.1 and 18 distinct time points (up to Day 60) for GIST-430 xenograft model. A longitudinal model based on the generalized estimating equations approach (Gaussian model with identity link and an autoregressive correlation structure) was used to model the effect of treatment and time on (the logarithm of) tumor volume. A linear time-effect was included in the model for the logarithm of tumor volume and interacted with treatment. Tumor volumes were compared between treatment groups using the two-sided Mann–Whitney test with a type I error of 5%. The package geepack was used for these computations (27).

## Results

### IM in combination with MK-4440 has enhanced viability on in vitro GIST cell growth

We have previously shown that targeting the KIT signaling pathway vertically at different nodes (i.e. at KIT using IM, in combination with downstream inhibition of AKT) demonstrated significant efficacy in an IM-sensitive GIST model. Here we sought to determine if this “vertical” inhibition could be an effective strategy in less IM-sensitive models. We selected MK-4440, a pan-AKT allosteric inhibitor, for these combination studies.

We first evaluated the effects of MK-4440 and IM on the growth of a panel of GIST cell lines: GIST-T1 (IM-sensitive), GIST-T1/829 (IM-resistant), and GIST430 (IM-resistant), as single agents and in combination at increasing molar ratios. **Figures 1A, B and C** show single agent dose response curves for GIST-T1, GIST-T1/829 and GIST430, respectively. We first estimated the LD50 for each agent in the GIST-T1 cell line (**Figures 1A, left panels**) and the LD30 for each agent in the GIST-T1/829 and GIST430 cell lines (**Figures 1B and C, left panels**). We then treated each line with increasing doses of the two drugs in a fixed ratio at their LD50 (GIST-T1) or LD30s (GIST430, GIST-T1/829) (**Figures 1A, B** and **C, third panel**). To quantify synergy, combination index (CI) values were calculated (**Figures 1A, B** and **C, last panel:** CI values <1 are considered synergistic). The CI_LD50_ value for GIST-T1 was 0.171. The CI_LD30_ values for GIST430 and GIST-T1/829 were 0.079 and 0.0595, respectively, indicating possible synergy in all three lines. These synergies were then established to be significant via a bootstrap statistic (28).

**Figure 1.**
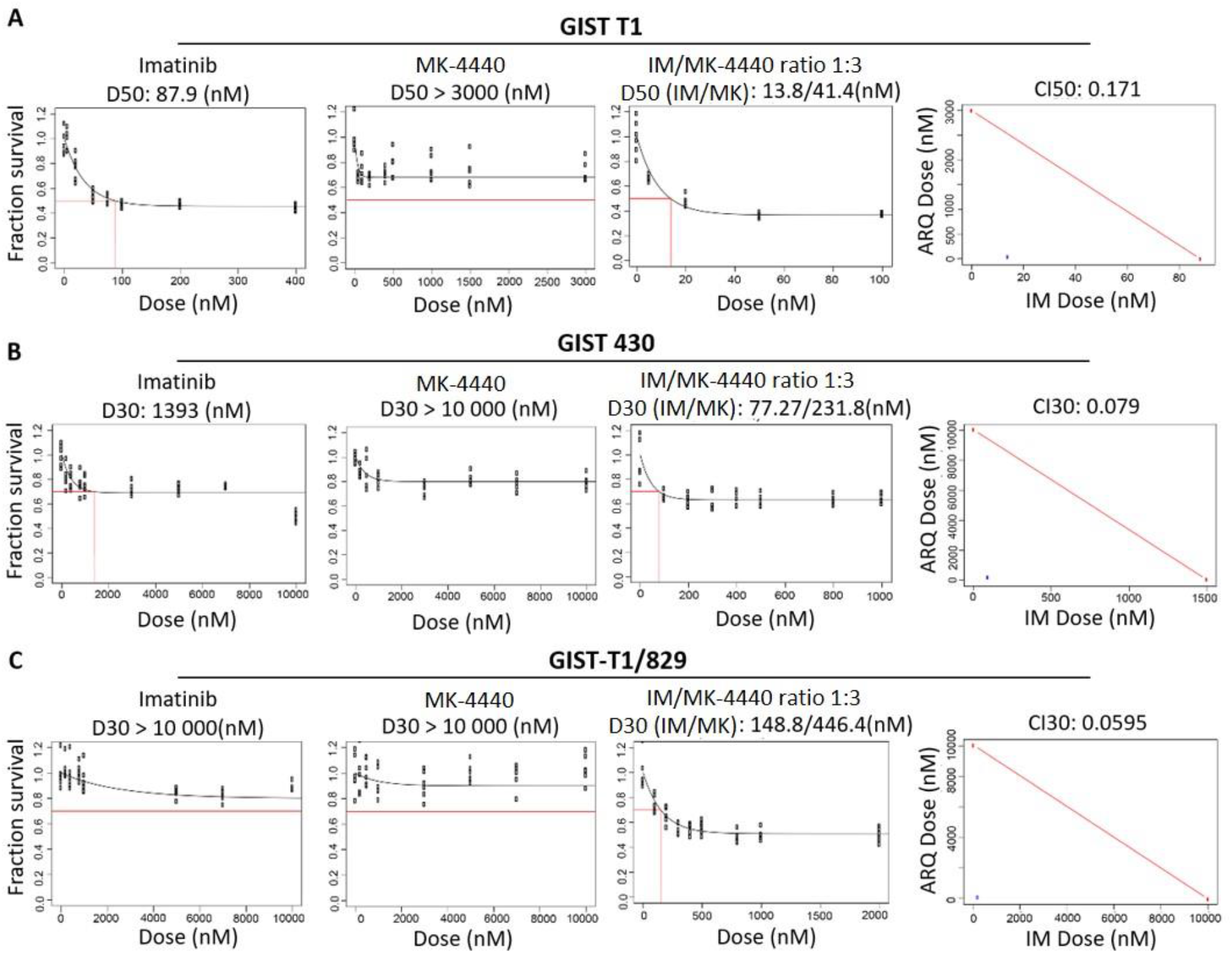
MK-4440 and IM have enhanced combination on in vitro GIST cell growth. Panels 1 & 2, Dose response curves for single agents (IM, MK-4440) in GIST-T1 **(A)**, GIST430 **(B)** and GIST-T1/829 **(C)** cell lines. Values represent the mean of survival compared to control cells (n=6). **Panel 3**, Dose response curve representing increasing series of combinations in GIST-T1 **(A)** GIST430 **(B)** and GIST-T1/829 **(C)** cell lines. Red box indicates estimation of LD50 (GIST-T1, **A**) or LD30 (GIST430, **B** and GIST-T1/829, **C**) concentration for combination of drugs. **Panel 4**, Single point (blue) on isobole curve for 50% kill (GIST-T1, **A**) or 30% kill (GIST430, **B** and GIST-T1/829, **C**). Red line indicates 50% (GIST-T1) isobole or 30% isobole for GIST430 and GIST-T1/829 for strictly additive effect. CI_LD50_ in GIST-T1 is 0.171 and CI_LD30_ in GIST430 and GIST-T1/829 are 0.079 and 0.0595, respectively, all found within the synergistic triangle.

### Combination treatment inhibits KIT signaling and increases PDCD4 expression

To assess the effects of drug treatment on signaling downstream of KIT, we performed immunoblotting on the same panel of GIST cell lines treated with IM, MK-4440, or the combination (**Figure 2**). As expected, IM at 40nM significantly reduced activation of KIT and downstream effectors, MAPK and AKT in the IM-sensitive cell line, GIST-T1. In IM-resistant lines GIST-T1/829 and GIST430, phosphorylation of AKT and downstream kinase, S6 ribosomal protein were less affected by IM treatment at the higher dose of 1μM. Inhibition of AKT was observed in all four cell lines treated with MK-4440. Interestingly, a significant decrease in the activation of S6 was observed in the GIST-T1 combination-treated cells compared to cells treated with single agents.

**Figure 2.**
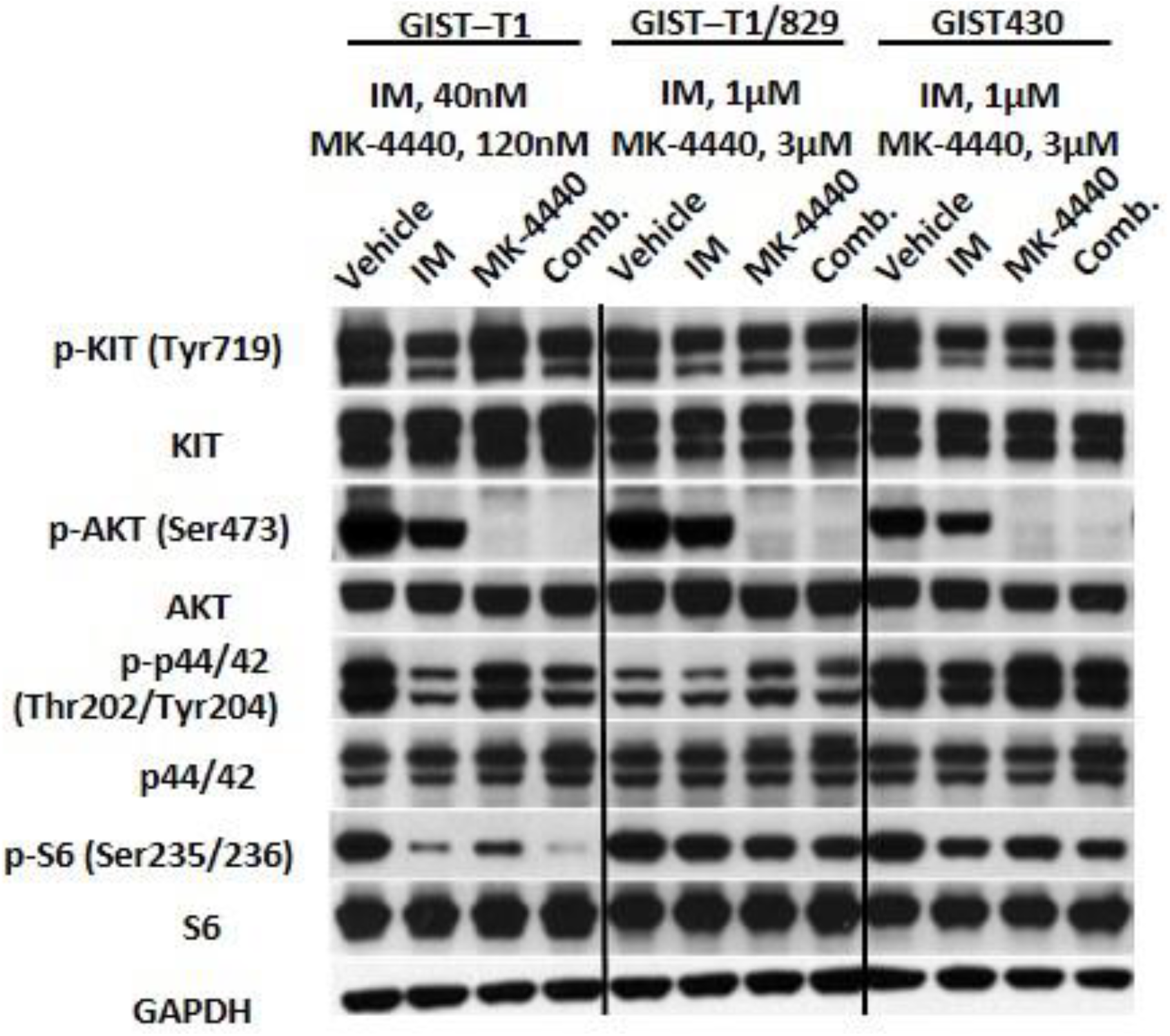
The combination of IM and MK-4440 inhibits constitutive activation of effectors downstream of KIT. Immunoblot assays of WCEs from GIST-T1 treated with 40nM IM, 120nM MK-4440 or the combination; and GIST430 and GIST-T1/829 treated with 1 μM IM, 3 μM MK-4440, or the combination. All treatments were twenty hours in duration. Equal quantities of WCE from each sample were subjected to immunoblotting with specific antibodies, as indicated. GAPDH served as loading control.

Next, we treated the isogenic cell lines, GIST-T1 and GIST-T1/829, with the combination of IM + MK-4440 or vehicle for 20 hours at the same doses described above and subjected them to LC-MS/MS proteomic analysis in order to understand the mechanism of the combination’s increased efficacy. A total of 3,120 unique proteins were identified in at least one experimental condition in all three replicates in the GIST-T1/829 group, and 2,837 unique proteins were identified in at least one experimental condition in all three replicates in the GIST-T1 group. From this complete list, we identified 40 downregulated and 13 upregulated proteins following combination treatment in GIST-T1 cells, while 7 downregulated and 5 upregulated proteins were found in GIST-T1/829 cells **(Table 1, Data file S2)**. Volcano plot analysis of fold change label free quantification (LFQ) values revealed proteins exhibiting differential protein abundance in GIST-T1 (**Figure 3A**) and GIST-T1/829 (**Figure 3B**) following treatment with IM+ MK-4440 combination. Interestingly, the IM-resistant GIST-T1/829 cells had considerably fewer differentially expressed proteins compared to their IM-sensitive counterpart, GIST-T1. In addition, groups of upregulated and downregulated proteins between GIST-T1 and GIST-T1/829 lines were very dissimilar. Only two downregulated proteins, ODC1 and TCAF1, were shared between the two cell lines (**Figure 3C)** and 2 upregulated proteins, PDCD4 and ATP5D, were common between GIST-T1 and GIST-T1/829 (**Figure 3D)**.

**Table 1.** Differentially expressed proteins upon treatment of GIST cells with IM/MK-4440 combination for 20h.

|  |  |  |  |  |  |
| --- | --- | --- | --- | --- | --- |
| GIST-T1 |  |  |  | GIST-T1/829 |  |
| Significantly downregulated upon treatment |  |  |  |  |  |
| ABCF2 | FADS2 | MKI67 | SQLE | ASAH1 |  |
| ACACA | FBN1 | NAV1 | SQSTM1 | ODC1 |  |
| AGFG1 | HMGCS1 | ODC1 | TACC3 | PTGS2 |  |
| ARMC9 | HN1 | PFN2 | TAX1BP3 | RAC2 |  |
| ATAD2 | HSBP1 | PRKCI | TCAF1 | RPL35A |  |
| BMP2K | IRF2BP2 | PRRC2A | TGFB1I1 | SPAG5 |  |
| CASP7 | KIF11 | RRM2 | TPX2 | TCAF1 |  |
| CBX1 | KPNA2 | SEC62 | TYMS |  |  |
| EIF1 | LSM14B | SLC4A1AP | ZFYVE16 |  |  |
| EMC8 | MAGT1 | SPRY4 | ZPR1 |  |  |
| Significantly upregulated upon treatment |  |  |  |  |  |
| ATP5D | H1FX | MRPL22 | PBXIP1 | ATP5D | NDUFAB1 |
| BANF1 | HIST1H1C | NDUFA4 | PDCD4 | HMGCR | PDCD4 |
| DERL1 | HIST1H4A | NDUFS4 | TMEM109 | ITM2C |  |
| GSTM2 |  |  |  |  |  |

**Figure 3.**
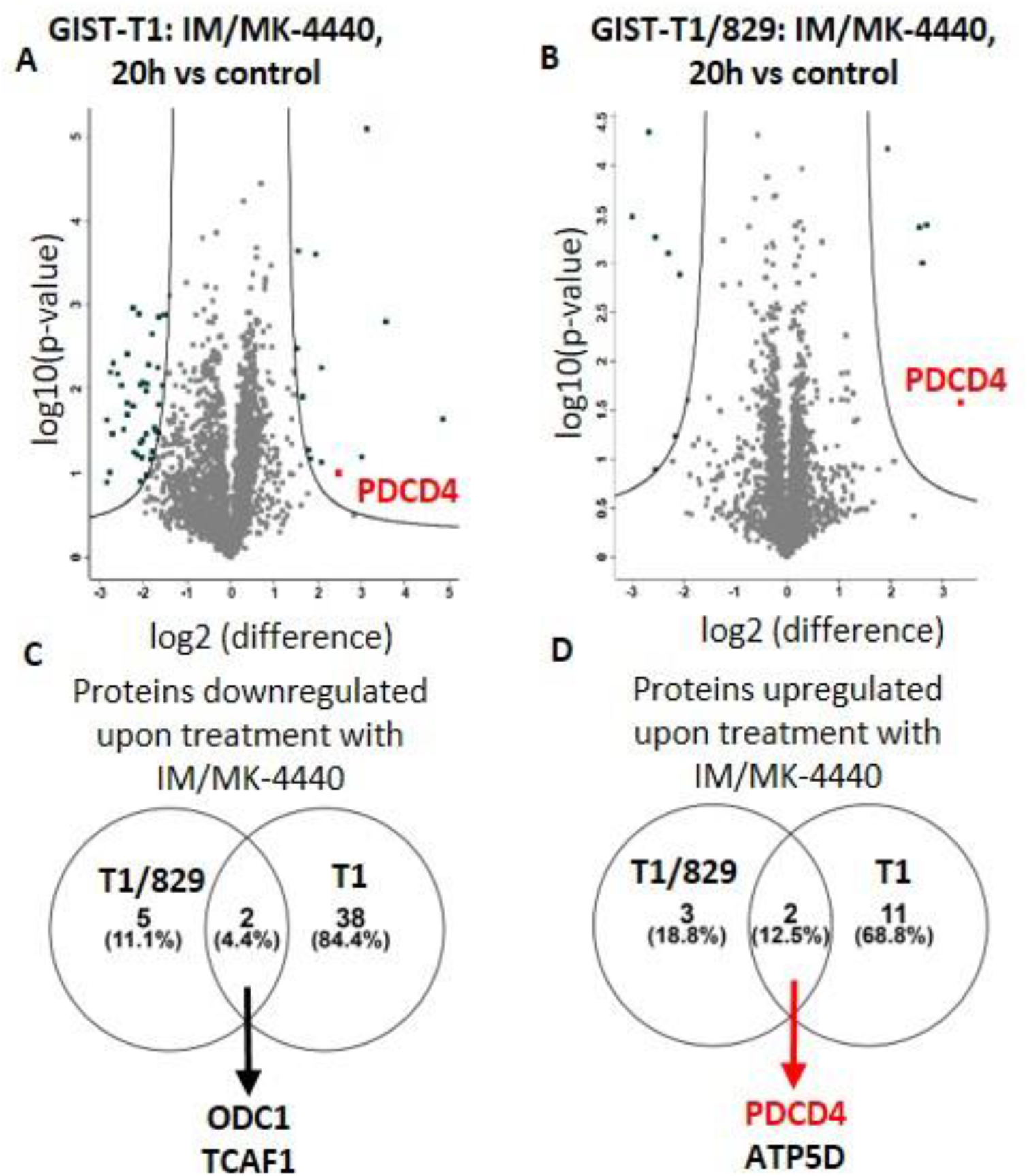
Total proteome changes upon IM/MK-4440 treatment. Volcano plot comparisons of total proteome in GIST-T1 (**A)** and GIST-T1/829 (**B)** cells treated with IM/MK-4440 combination versus vehicle (control) for 20h. Differences in log2 LFQ intensities among cell lysates from three experiments determined by paired t-test at FDR of <0.05 using Perseus software. Venn diagram comparisons of downregulated (**C)** and upregulated **(D)** proteins between GIST-T1and GIST-T1/829.

### Combination treatment causes cell cycle arrest and increases cell death

Programmed Cell Death 4 (PDCD4) is a tumor suppressor and apoptotic activator whose expression has been shown to be upregulated following PI3K/AKT inhibition leading to subsequent elevation of cell cycle inhibitor p27, in several malignancies. In order to evaluate the role of PDCD4, we performed immunoblotting on GIST cell lines treated with IM, MK-4440, or the combination **(Figure 4A)**. Combination treatment significantly increased PDCD4 expression, and, as expected, increased cleaved-PARP in all three GIST cell lines, suggesting increased apoptosis compared to cells treated with either single agent. Furthermore, elevated levels of p27 were observed in combination-treated GIST-T1 and GIST430, but not in GIST-T1/829 cells.

**Figure 4.**
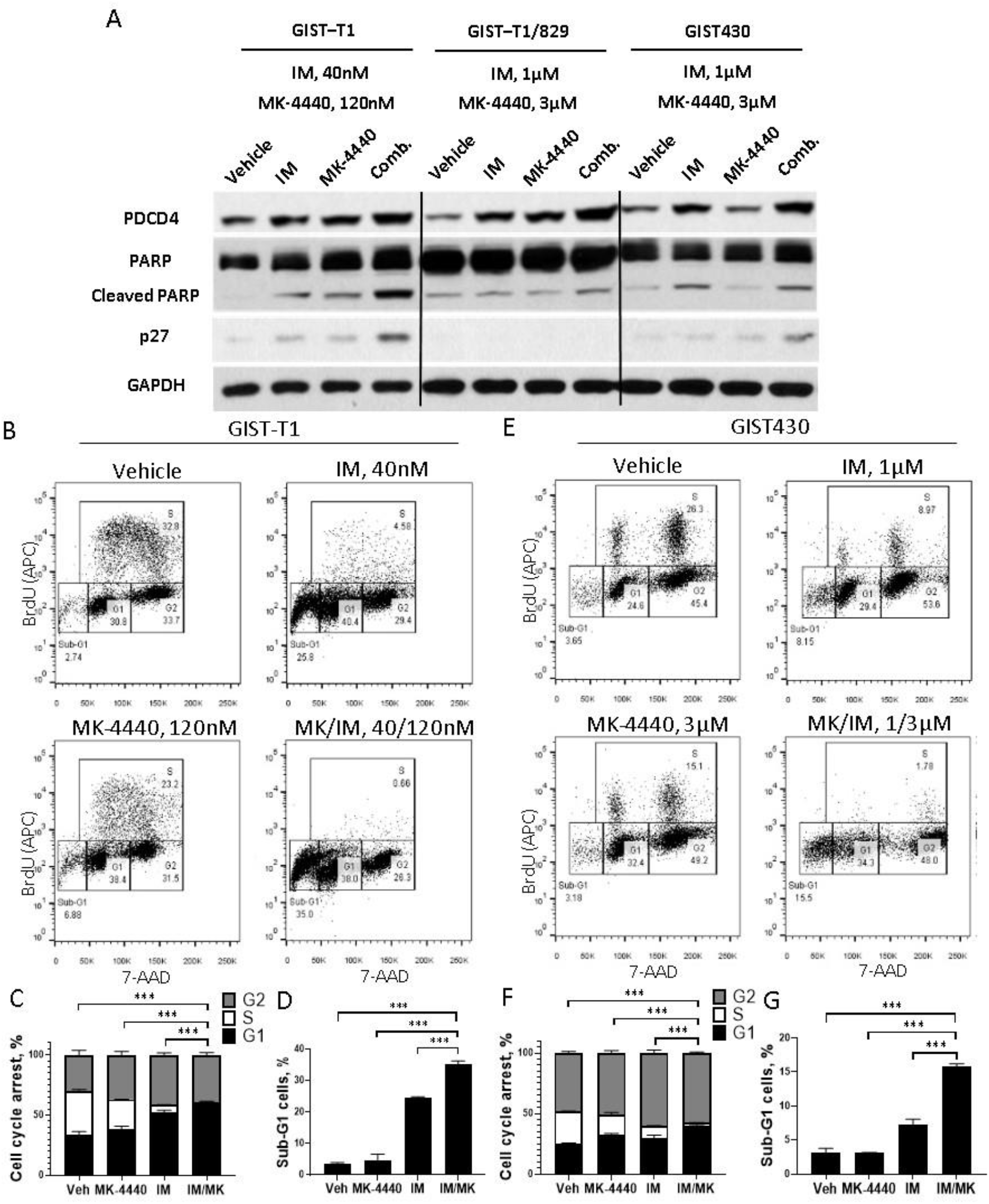
IM and MK-4440 combination induces cell cycle arrest and apoptosis. Immunoblot assays of WCEs from GIST-T1 treated with 40nM IM, 120nM MK-4440 or the combination; and GIST-T1/829 and GIST430 treated with 1 μM IM, 3 μM MK-4440, or the combination. All treatments were twenty hours in duration. Equal quantities of WCE from each sample were subjected to immunoblotting with specific antibodies, as indicated. GAPDH served as loading control (**A**). Representative flow cytometry plots (**B**) and quantification (**C, D**) of BrdU incorporation in GIST-T1 treated with 40nM IM, 120nM MK-4440 and combination for 72h. For comparisons between the combination (IM/ MK-4440) treatment vs. all monotherapies (Vehicle, IM, MK-4440) statistically significant reduction in S phase, increase in cell death (*** p < 0.00007), and G1 arrest (p < 0.009) were observed. Representative flow cytometry plots (**E**) and quantification (**F, G**) of BrdU incorporation in GIST430 treated with 1 μM IM, 3 μM MK-4440 and combination for 72h. For comparisons between the combination (IM/ MK-4440) treatment vs. all monotherapies (Vehicle, IM, MK-4440) statistically significant reduction in S phase, increase in cell death (*** p < 0.0002), and G1 arrest (p < 0.002) were observed. Data represent mean of three experiments + SD.

The effect of pharmacological inhibition of KIT and AKT on cell cycle dynamics in GIST cells was measured with a BrdU assay. GIST-T1 and GIST430 cells treated with vehicle, IM, MK-4440 or the combination, were analyzed by flow cytometry after BrdU incorporation and subsequent antibody binding in combination with direct 7-AAD staining **(Figure 4B, E)**. IM treatment induced G2 arrest in GIST430 cells, and G1/G2 arrest in GIST-T1 cells, whereas MK-4440 treatment alone caused very minor changes to cell cycle dynamics compared to vehicle. Combination treatment induced both G1 and G2-phase cell cycle arrest with significantly fewer cells in S-phase, compared to cells treated with either monotherapy **(Figure 4C, D)**. In addition, cells treated with the combination exhibited a 1.5-fold increase in the sub-G1 population in both GIST-T1 and GIST430 cells, indicating increased cell death compared to IM monotherapy treatment **(Figure 4F, G)**.

### Combination treatment reduces tumor growth *in vivo* in IM-sensitive GIST only

Based on strong *in vitro* data showing a synergistic relationship between IM and MK-4440 (**Figure 1)** and effects of the combination on cell cycle and apoptosis (**Figure 4)** in both GIST-T1 and GIST430 cell lines, we next sought to evaluate this combination *in vivo*. For these studies we used IM-sensitive GIST-T1 xenografts, IM-resistant GIST430 xenografts modeling acquired secondary drug resistance, as well as a patient-derived GIST xenograft (PDX) established by our group, PDX9.1. PDX9.1 has been serially passaged for 8 passages, and has maintained the basic histological, immunohistochemical, and genetic characteristics of the patient tumor (**Figure 5)**. This PDX harbors a KIT mutant isoform (exon 9, p.A502_Y503dup) modeling GIST requiring higher dose IM therapy.

**Figure 5.**
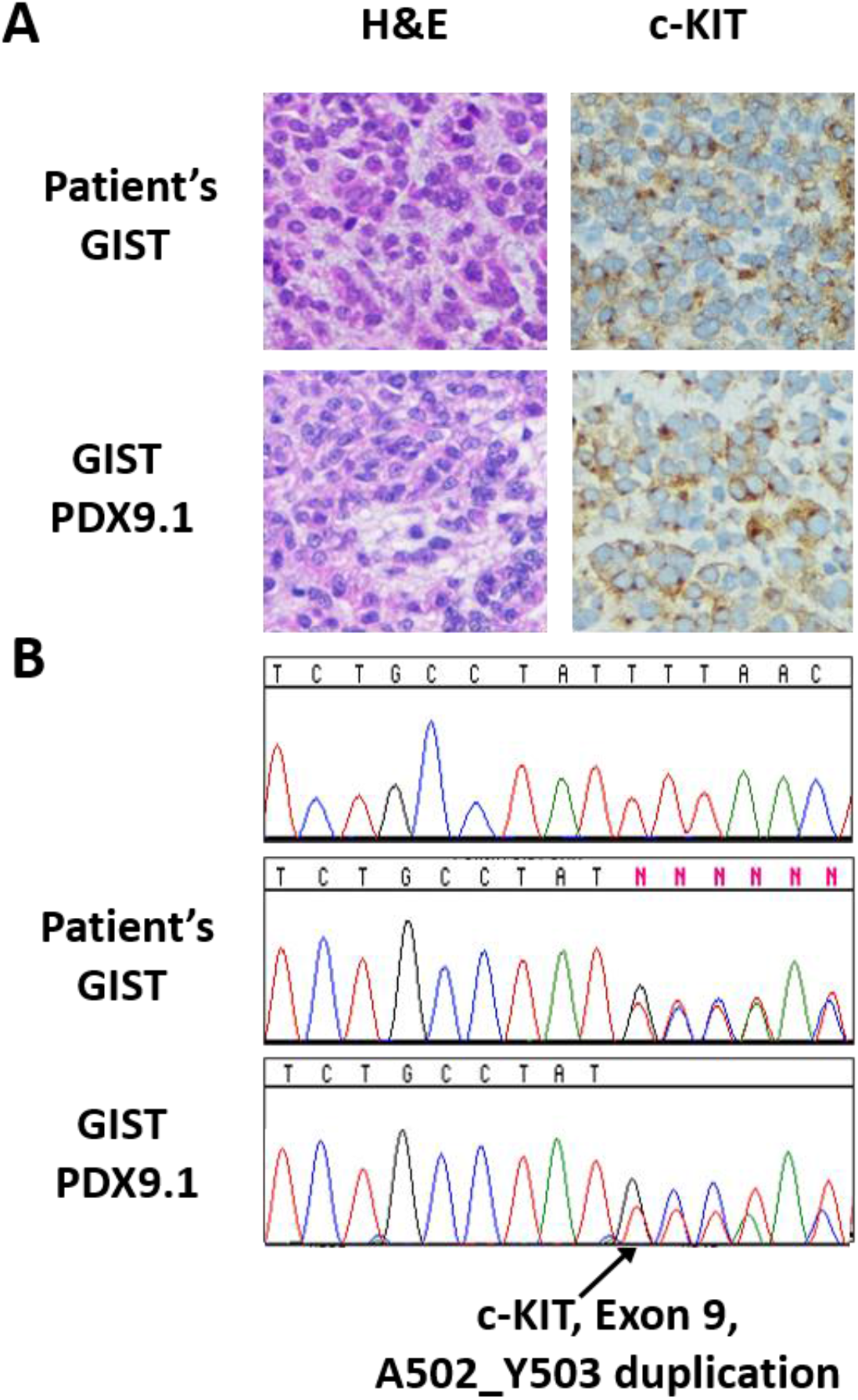
Immunohistochemical and mutational analysis of a primary GIST and matched PDX model. **(A)** H&E and c-KIT (CD117) staining for primary GIST (top) and matched GIST PDX9.1 (bottom). **(B)** Sanger sequencing analysis identified the A502_Y503 duplication in exon 9 of *KIT* in both primary GIST (middle) and matched GIST PDX9.1 (bottom), but not in a non-tumor specimen (top).

Xenografts were established subcutaneously in a total of 7 to 10 mice per cell line and randomized into four treatment arms: arm 1: vehicle, arm 2: IM, arm 3: MK-4440 and, arm 4: IM/ MK-4440 combination. Impressively, GIST-T1 xenografts treated with the combination showed significantly greater disease stabilization and, in some mice, tumor regression compared to the standard of care, IM (p < 0.0006) (**Figure 6)**. However, in both IM-resistant GIST models, GIST430 and PDX9.1, no statistically significant difference was observed in combination treated mice compared to IM, although trends indicating improvement over monotherapies were observed (**Figure 6**).

**Figure 6.**
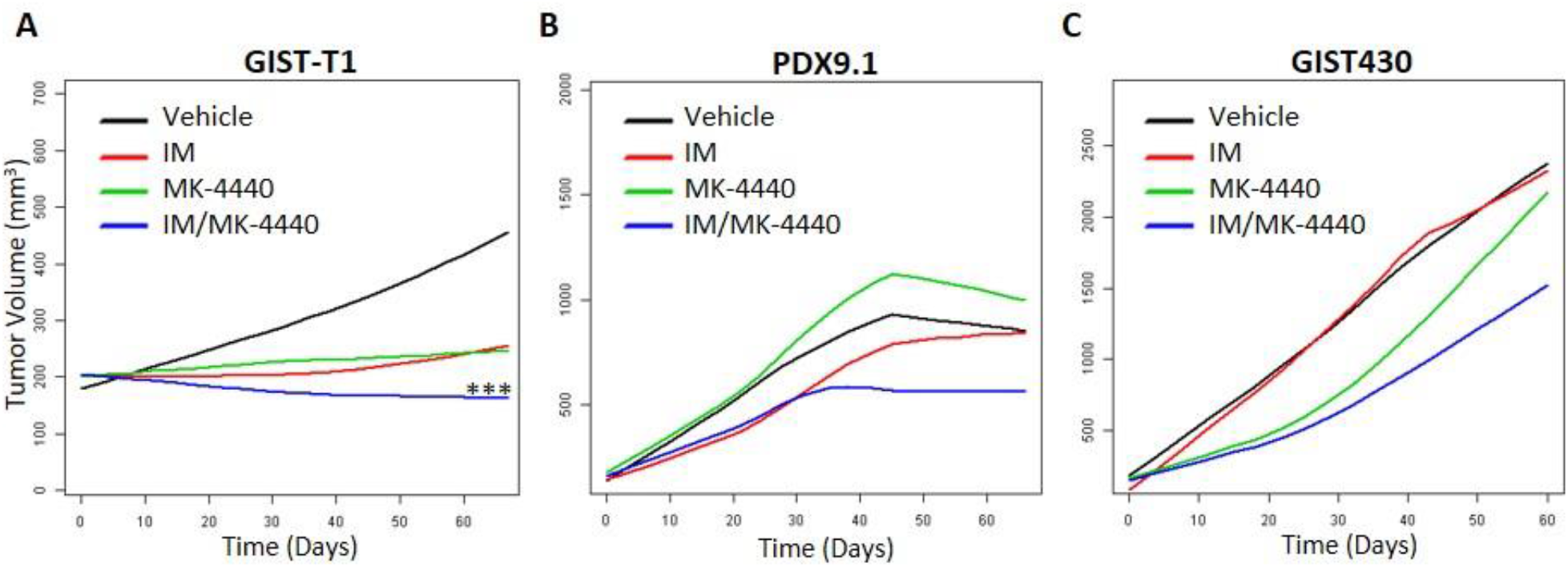
Combination of IM and MK-4440 inhibits GIST growth in vivo. Smoothed tumor growth curves (tumor volume vs. time) were computed for each treatment using the lowess smoother in the R statistical language. **(A)** Statistically significant decrease in the rate of GIST-T1 xenograft tumor growth was observed for IM/MK-4440 combination (blue) vs. standard monotherapy with IM (red) (p < 0.0006). Tendency to decrease in the rate of tumor growth in PDX9.1 **(B)** and GIST430 xenograft **(C)** models under treatment with IM/MK-4440 combination (blue) were observed with no statistical significance in comparison with IM monotherapy (red).

## Discussion

For the last twenty years, IM has remained the first-line therapeutic option for the majority of GIST. During this time, three additional RTK inhibitors (sunitinib, regorafenib, ripretinib) have been approved for treatment of IM-refractory tumors; however, progression free survival for all of these agents is measured in months. Once all approved lines of therapy are ineffective, patients with advanced GIST are left without further treatment options. Therefore, abrogating or delaying development of acquired resistance to IM could have potential clinical benefit. Activation of the PI3K/AKT pathway has been shown to predict and promote resistance to IM in GIST (19, 21, 22, 29). Several preclinical studies have reported significant efficacy of IM in combination with PI3K inhibitors in IM-sensitive GIST models; however, mixed reports regarding efficacy of this combination have been reported in various IM-resistant models (30-33). Previously, our group had shown treatment with IM in combination with the pan-AKT inhibitor MK-2206 demonstrated increased efficacy in IM-sensitive and resistant GIST cell lines, as well as, extended disease stabilization and improved survival in an IM-sensitive xenograft model compared to IM treatment alone (19).

In this study we sought to determine if AKT inhibition in combination with IM could be an effective strategy in less IM-sensitive models, including an established PDX GIST model which possesses an exon 9 KIT mutation. We selected MK-4440, a pan-AKT allosteric inhibitor, as allosteric inhibitors of AKT have been shown to more potently induce cell death in wild-type AKT cell lines compared to ATP-competitive inhibitors (34). MK-4440 is currently being tested in a phase 1 clinical trial as a single agent or in combination with other anti-cancer agents in solid tumors (NCT02761694). We demonstrated increased efficacy between MK-4440 and IM at a 3:1 ratio in both IM-sensitive and resistant cell lines, although IC50s were not achieved for either of the resistant lines due to inherent IM resistance. To gain insight into the mechanism(s) responsible for the superior efficacy of this combination, we performed proteomic analysis using isogenic IM-sensitive and resistant GIST cell lines. This analysis focused our attention on Programmed Cell Death 4 (PDCD4), whose expression was significantly upregulated in combination treated IM-sensitive and resistant cell lines compared to either agent alone.

PDCD4 is known for its extensive role as a tumor suppressor and apoptosis activator. Loss of PDCD4 has been associated with tumor transformation and poor prognosis in a variety of cancers (35). Interestingly, loss or downregulation of PDCD4 expression has been observed in 68% of GIST and is associated with tumor progression (36). pS6 has been shown to phosphorylate PDCD4 leading to its degradation (37, 38). In both our previous study (19) and in this work, the combination of IM and AKT inhibition (MK-2206 and MK-4440) led to a decrease in S6 phosphorylation compared to either single agent. Previous studies in AML, endometrial and ovarian cancer, showed inhibition of PI3K/AKT led to subsequent upregulation of PDCD4 and concomitant upregulation of cell cycle inhibitor, p27 (39-41). Combination treatment in our panel of IM-sensitive and resistant GIST cell lines demonstrated elevated PDCD4, p27 expression, and increased cleaved-PARP. Additionally, mass spectrometry data revealed downregulation of ODC1 protein in both IM-sensitive and resistant isogenic cell lines. ODC1 is involved in DNA synthesis, peaking twice during the cell cycle, at the G1/S and at G2/M transitions, therefore downregulation of ODC1 confirms cell cycle arrest and lack of S/M phase cells upon treatment with IM/ MK-4440 (42-44). BrdU assays indicated combination treatment in both IM-sensitive and resistant cell lines induced both G1 and G2-phase cell cycle arrest with significantly fewer cells in S-phase as well as increased sub-G1 population compared to either single agent. Taken together, these results demonstrate that dual inhibition of KIT and AKT in both IM-sensitive and resistant GIST cells leads to decreased activation of pS6, associated PDCD4 upregulation, cell cycle arrest through upregulation of p27, and increased apoptosis (**Figure 7)**.

**Figure 7.**
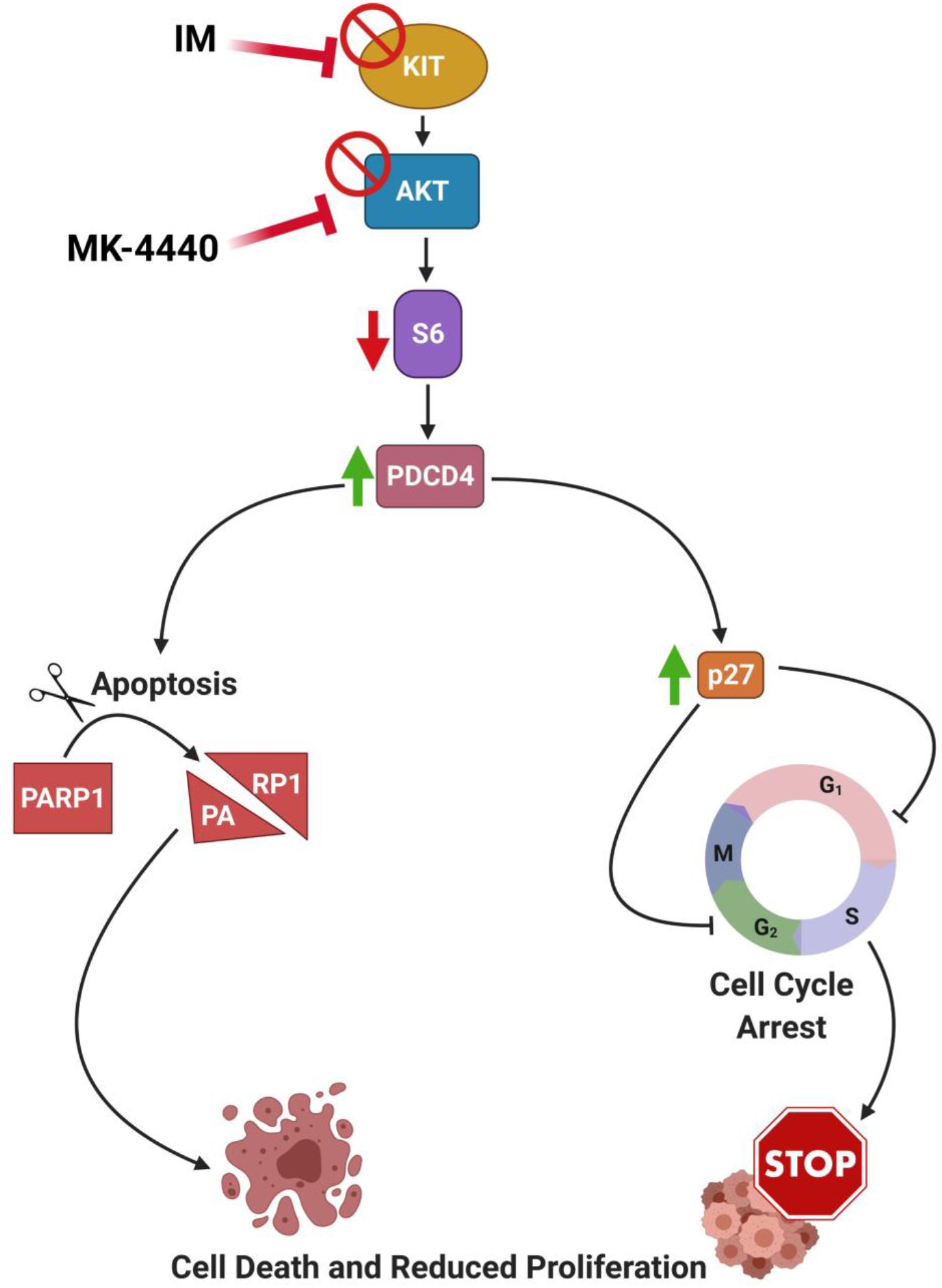
Proposed mechanism of dual inhibition of KIT and AKT in GIST. Dual inhibition of KIT and AKT in GIST cells leads to decreased activation of pS6, associated PDCD4 upregulation, cell cycle arrest through upregulation of p27, and increased apoptosis.

The results of these *in vitro* studies provided justification for evaluating such an approach *in vivo* to determine whether this combination would improve efficacy of IM, increase time to resistance in a sensitive model, and/or have potential in the resistance setting. Compared to standard of care IM, combination treated GIST-T1 xenografts (IM-sensitive) displayed significantly greater disease stabilization and in some instances, tumor regression. However, in IM-resistant models, GIST430 and PDX9.1, no statistically significant differences occurred, although trends suggesting superior effect of combination compared to monotherapies were observed.

While our *in vitro* combination drug studies generated significant synergistic CI values in the IM-resistant cell lines, the *in vivo* experiments instead exhibited trends towards superior efficacy in the IM-resistant mouse models that were, however, not statistically significant. High concentrations of both IM and MK-4440 were utilized in the IM-resistant cell line cytotoxicity, proteomic and BrdU assays, but dose limiting toxicities in the animal models precluded the use of increased drug dosing. Future studies evaluating PI3K/AKT inhibition in the IM-resistant setting should evaluate its combination with RTK inhibitors with alternative mechanisms of action, such as ripretinib (switch-pocket inhibitor) or avapritinib (type 1 inhibitor) which should have greater efficacy as single agents compared to IM. Interestingly, AKT inhibition has recently been reported to enhance cytotoxicity of topoisomerase II inhibitors in GIST (45) suggesting an additional option for potential combination. In conclusion, AKT inhibition in combination with IM demonstrated significant, lasting efficacy in IM-sensitive GIST, indicating justification for the development of future clinical trials evaluating this combination in primary GIST. In the resistant setting, however, alternative agents will need to be tested in combination with AKT inhibition.

## Acknowledgements

This work was supported by NIH CORE grant CA06927 (FCCC), NCI R00 CA158065 (L.R), NCI R01 CA212662 (L.R.) and WJ Avery Fellowship (S.Y.). We would like to acknowledge the following facilities at FCCC for their work contributing to this manuscript: Cell Sorting, High Throughput Screening, Genotyping and Real-Time PCR, and the Laboratory Animal Facility. The authors would especially like to thank the David Foundation and the GIST Cancer Research Fund for their continued support.

