## Supplemental Figure 1 for "Combined Inhibition of AKT and KIT Restores Expression of Programmed Cell Death 4 (PDCD4) in Gastrointestinal Stromal Tumor"

**Protein Extraction and LC-MS/MS Analysis**

GIST cell lines were treated for 20 hours with IM/MK-4440 combination at the following concentrations, 40nM/120nM (GIST-T), 1µM/3µM (GIST-T1/829) or corresponding amount of DMSO (control) in three biological replicates. Cells were lysed in 8M urea, 50 mM ammonium bicarbonate (ABC) and benzonase 24U/100mL with vigorous shaking (20 Hz for 10 min at room temperature by Retsch MM301 instrument). Lysate was centrifuged at 14,000xg for 10 minutes to remove cellular debris, and protein concentration of the supernatant was then determined using bicinchoninic acid (BCA) protein assay (Thermo Scientific). Protein disulfide bonds were reduced with 5mM tris(2-carboxyethyl)phosphine (TCEP) at 30°C for 60 min, and cysteines were subsequently alkylated with 15 mM iodoacetamide (IAA) in the dark at room temperature for 30 min. Urea was then diluted to 1 M urea using 50 mM ABC, and proteins were subjected to overnight digestion with mass spec grade Trypsin/Lys-C mix (Promega). Following digestion, samples were acidified with formic acid (FA) and subsequently desalted using AssayMap C18 cartridges mounted on an Agilent AssayMap BRAVO liquid handling system. Cartridges were first conditioned with 100% acetonitrile (ACN) followed by 0.1% FA, samples were then loaded, washed with 0.1% FA, and peptides eluted with 60% ACN, 0.1% FA. Finally, the organic solvent was diluted to allow peptide quantification using a NanoDrop spectrophotometer (Thermo Scientific), and then the remaining of the sample was dried in a SpeedVac concentrator.

Dried peptide fractions were reconstituted with 2% ACN, 0.1% FA and analyzed by LC-MS/MS using a Proxeon EASY nanoLC system (Thermo Fisher Scientific) coupled to an Orbitrap Fusion Lumos mass spectrometer (Thermo Fisher Scientific). Peptides were separated using an analytical C18 Aurora column (75µm x 250 mm, 1.6µm particles; IonOpticks) at a flow rate of 300 nL/min using a 75-min gradient: 1% to 6% B in 1 min, 6% to 23% B in 44 min, 23% to 34% B in 28 min, and 27% to 48% B in 2 min (A= FA 0.1%; B=80% ACN: 0.1% FA). The mass spectrometer was operated in positive data-dependent acquisition mode. MS1 spectra were measured in the Orbitrap with a resolution of 60,000, at accumulation gain control (AGC) target of 4e5 with maximum injection time of 50 ms, and within a mass range from 375 to 1500 m/z. The instrument was set to run in top speed mode with 1-second cycles for the survey and the MS/MS scans. After a survey scan, tandem MS was performed on the most abundant precursors with charge state between +2 and +7 by isolating them in the quadrupole with an isolation window of 0.7 m/z. Precursors were fragmented with higher-energy collisional dissociation (HCD) with normalized collision energy of 30% and the resulting fragments were detected in the ion trap in rapid scan mode at AGC of 1e4 and maximum injection time of 35 ms. The dynamic exclusion was set to 20 sec with a 10 ppm mass tolerance around the precursor.

All mass spectrometry proteomics data including the raw data (".RAW") and search result files from MaxQuant have been deposited to ProteomeXchange Consortium via MASSIVE partner repository with the dataset identifier PXD023717.

**Data Processing and Analysis**

All mass spectra were analyzed with MaxQuant software version 1.5.5.1. MS/MS spectra and searched against the curated (Swiss-Prot) Homo sapiens Uniprot protein sequence database (downloaded in January 2019) and GPM cRAP sequences (commonly known protein contaminants). Precursor mass tolerance was set to 20ppm and 4.5ppm for the first search where initial mass recalibration was completed and for the main search, respectively. Product ions were searched with a mass tolerance 0.5 Da. The maximum precursor ion charge state used for searching was 7. Carbamidomethylation of cysteine was searched as a fixed modification, while oxidation of methionine and acetylation of protein N-terminal were searched as variable modifications. Enzyme was set to trypsin in a specific mode and a maximum of two missed cleavages was allowed for searching. The target-decoy-based false discovery rate (FDR) filter for spectrum and protein identification was set to 1%.

Log2-transformed LFQ intensities for all 5407 detected proteins in four treatment subgroups (GIST-T1, GIST-T1+IM/MK-4440, GIST-T1/829, GIST-T1/829+IM/MK-4440,) in three biological replicates, including the potential contaminants, zero intensity proteins, as well as single-peptide identified proteins are contained in **Data file S1**. Common contaminants and proteins identified by site only were removed and only proteins identified by >1 unique peptides (4872proteins) were qualified for subsequent analysis using Perseus software (1.6.10.50) (24). Only proteins identified in all three biological replicates in at least one experimental condition (control or under IM/MK-4440 treatment) were further considered for statistical analysis (totally 3120 proteins in GIST-T1/829 cell line subgroup and 2837 proteins in GIST-T1 subgroup). In order to save normal distribution of the $Log_2$ transformed intensity histogram and to simulate signals from low abundant proteins, the filtered LFQ data were subjected to imputation of missing values performed from the normal distribution (width 0.3 and down shift 1.8) (25). After normalization, Pearson's Correlation was used to identify potential outliers, as a result there were no replicates removed from the analysis.

Two-sided Student’s t-tests were performed to identify significantly changing proteins among control and IM/MK-4440 treated cells. *P*-values were adjusted for using permutation-based false-discovery rate (FDR = 0.05, s0 = 2, and 250 randomizations). **Data file S2** contains lists proteins qualified for the analysis, as well as significance test results. Volcano plots depicting Log2 fold change differences in protein abundance following IM/MK-4440 treatment was generated with Perseus for each cell line. Proteins significantly up- and downregulated upon IM/MK-4440 treatment were compared between GIST-T1 and GIST-T1/829 cell lines with Venny 2.1 plotter.
